# Coverage-based detection of copy number alterations in mixed samples using DNA sequencing data: a theoretical framework for evaluating statistical power

**DOI:** 10.1101/413690

**Authors:** Shweta Ramdas, Yanchao Pan, Jun Z Li

## Abstract

DNA sequencing can discover not only single-base variants but also copy-number alterations (CNAs). In shotgun sequencing, regions of CNAs show step-wise changes in read depth when compared to adjacent “normal” regions, allowing their detection by parametric statistical tests that compare the mean coverage in suspected regions against that of a baseline distribution. Traditionally, the power of such a test depends on (1) the integer number of copy number change, (2) the overall sequencing depth, (3) the length of the CNA region, (4) the read length and (5) the variation of coverage along the genome, which depends on many experimental factors, including whether the chosen platform is whole-genome, whole-exome, or targeted-panel sequencing. In cases involving inadvertent sample mixing or genuine somatic mosaicism, power also depends on the mixing ratio. However, the analysis of statistical power that considers the interplay of all these factors has not been systematically developed. Here we present a general analytical framework and a series of simulations that explore situations from the simplest to the increasingly multifactorial. Specifically, we expand the expression of power to include not just the known factors but also one or both of two complications: (1) the dispersion of read depth around the mean beyond the independent sampling-by-sequencing assumption, and (2) the reduced fraction of the CNA-bearing sample (“purity”) as seen in studies of intratumor heterogeneity or in clinical monitoring of minimal residual disease. We describe the analytical formula and their simplifications in special cases, and share the extendable scripts for others to perform customized power analysis using study-specific parameters. As study designs vary and technologies continue to evolve, the input data and the noise characteristics will change depending on the practical situation. We present two use cases commonly encountered in cancer research: ultra-shallow whole-genome sequencing for detecting large, chromosome-scale events, and targeted ultra-deep sequencing for surveillance of known CNAs in rare tumor clones in the task of sensitive detection of cancer relapse or metastasis. We also present an online calculator at https://shiny.med.umich.edu/apps/hanyou/CNV_Detection_Power_Calculator/.

## 2 Background

A common approach for CNA-detection using genomic data relies on the telltale stepwise change of apparent DNA “dosage” in a genomic region (Olshen et al., 2004). For data from microarray-based comparative genomic hybridization (arrayCGH) or single nucleotide polymorphism (SNP) genotyping, the raw signal for DNA dosage is the total hybridization intensity at the individual *locus*, which is either the arrayCGH probe location or the SNP site. The spatial distribution of these measured loci along the genome has been determined by how the genetic markers were selected during the design stage of each array platform. A more recent, rapidly adopted technology is high-throughput DNA sequencing, where the observed number of mapped reads in a given genomic interval reflects the regional variation of DNA in the input material. In this study we mainly consider sequencing data, and do not make the distinction between germline DNA or somatic DNA. We will use CNA and CNV (copy number variation) interchangeably. When discussing sample purity, we refer to either the unintended mixing of *two* germline DNA samples or the coexistence of two somatic cell populations (such as cancer cell clones). We assume that all the align-able reads have been correctly mapped, thus sidestepping the situation when segmental duplications might cause some reads to be ambiguously mapped to multiple locations. We will not address the situation of copy-neutral structural variation such as translocations or inversions. Instead we focus on copy number changes.

For both microarray data and sequencing data, regional variation of signal intensity along the genome presents a major challenge for detecting step-wise changes. Systematic biases, such as those due to local GC-content, are technology-dependent and can be largely corrected by appropriate normalization steps based on local bias patterns learned from a large sample, for example (Diskin et al., 2008). In this study we will not dwell on the issue of bias correction, and will assume that the best procedures for correcting systematic variation will have been applied in early stages of data preparation. Instead we will examine the impact of signal over-dispersion, i.e., the “noise” of read depth along the genome beyond what is expected for random sampling from a Poisson distribution.

Read depth-based CNA detection can be formulated as a task of comparing of the mean between two“samples”, as implemented in parametric tests such as the Student’s t-test. Importantly, the *unit of observation* depends on the technology. For microarrays, the “data point” being sampled is the quantitative intensity of an arrayCGH probe or the total signal of the two alleles at a SNP site. Here the measurement units, and their spatial distribution, have been pre-defined by the chosen platform; and in general, the sampled locations tend to cover the genome at variable intervals. For whole-genome sequencing (WGS) data, the analyst can choose a window width and tally the total number of reads in consecutive windows, where each window provides a data point. If a CNA contains n data points, the task is to determine if the mean of such n observations is different from the mean of a null distribution, which can be established either with adjacent “normal” intervals or with a genome-wide aggregate of baseline regions.

We denote the haploid sequencing read depth as D (that is, for a dataset of 30X total coverage, the haploid coverage is 15), the length of the CNA to be detected as L, the size of the window into which markers/reads are binned as W, the ploidy of the CNA as N (normal diploid regions have N=2), and read length as l. We denote the purity of the sample as F, defined in this study as a two-way mixing of a normal sample at (1-F) and an aneuploidy sample at F. Currently, we do not consider three-way or more complex mixing.

If we ignore the reads falling on window boundaries, a window of length W would contain W/l reads laid end-to-end. At a depth of D this window is expected to contain X=2DW/l reads in a diploid region, and NDW/l reads in a region of ploidy N. When we use the mean (Avg(X)) to estimate the expected per-window read count 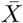, a t-like statistic can be constructed as the difference of the mean between the N-ploid CNA X_1_ and a diploid baseline X_2_, scaled by the standard error of this difference:

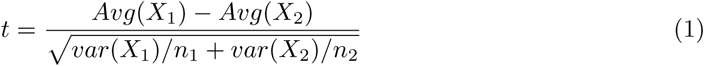

where *X*_1_ and *X*_2_ are the per-window read count in the CNA and outside the CNA, respectively. *n*_1_ and *n*_2_ are the number of independent measurements, i.e., the number of windows, in and outside the CNA, respectively. The numerator can be expressed as |N − 2|DW/l. A CNA of length L contains *n*_1_=L/W windows.

Next, we make the assumption that most of the genome is diploid and the baseline diploid regions can be pooled such that *n*_2_ is much larger than *n*_1_. By omitting the (1*/n*_2_) term in the denominator we can simplify Equation 1 to:

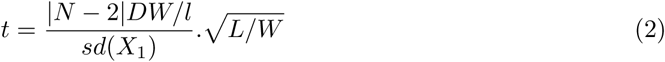

where the denominator is the standard deviation of X=NDW/l in the CNA region. The last term is 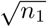, the square root of “sample size”, i.e., the number of windows in the N-ploid CNA.

We would like to re-emphasize that we use D to denote haploid read depth in order to make the expression less cluttered. For instance, in an experiment with a mean coverage of the diploid genome of 30X, D=15; and a region with a ploidy of 3 would be expected to have a coverage of 45X with an expected difference of D(N-2) of 15.

The t-like score in equations (1) and (2) represents the total signal in a power analysis, which will partition t into two components: *t* = *t*_*α*_ + *t*_*β*_, where *t*_*α*_ is the component for a significance level of *α*, and *t*_*β*_ is the component for a power of 1-*β*. In essence, the total t-like score is to be “spent” in two ways, one for a specific Type-1 error rate, and the rest for a specific Type-2 error rate. In the rest of the manuscript we will derive the expressions for the total t score, whereas in plotting the power curves we fix *t*_*α*_ for a per-event Type-1 error of 0.05. In real situations, the experiment-wide Type-1 error needs to be calculated with multiple-testing correction according to the number of CNAs to be detected for the entire DNA sample.

## 3 Motivation

As outlined above, a standard power analysis involves five key components: on the left side of the expression is the total weight of evidence, t, split into *t*_*α*_+*t*_*β*_, to account for the tradeoff between the false positive and false negative rates. On the right side of the expression is the effect size, divided by the standard deviation of this effect, and multiplied by the square root of the sample size. In the context of coverage-based CNA detection, the effect size is a function of four parameters: |N-2|DW/l, and sample size is L/W.

This study is motivated by the need to explore three factors that affect power. First, the choice of W brings two opposing consequences: a larger W increases effect size but reduces sample size. We will examine the net outcome of varying W. Second, the number of reads in consecutive windows may not vary according to a simple Poisson sampling process. Even after correcting for systematic regional biases, certain levels of over-dispersion may remain, leading to higher values in the denominator, and reduced power. We will introduce a variance inflation parameter in the power calculator. Third, sometimes the DNA sample represents the mixing of two cell populations (due to inadvertent sample mixing, tumor heterogeneity, donor-recipient mixing during transplantation, and maternal-fetal mixing in prenatal testing), and sometimes the CNA-bearing population is very rare. We denote the sample mixing ratio as F. Can a 10-fold reduction of “purity” be compensated by 10-fold deeper sequencing? We will examine questions like these, with or without the variance inflation factor.

The results below are organized into three parts. In #4 we examine the situation of 100% purity (F=1), first with and then without variance inflation. In #5 we examine the impact of sample impurity (F<1), again with and without inflation. In #6 we explore two use cases, one with very long CNAs (large L) but low sequencing depth (small D) and varying purity (large F); the other with high sequencing depth (large D) but low purity (small F) and varying L.

## 4 A single population (mixing ratio F=1)

### 4.1 No variance inflation (Poisson sampling)

In the simplest case, read count along the genome can be modeled as a Poisson sampling process where the variance of read counts across a string of consecutive windows equals the mean of these windows. The distribution of *X*_1_, the per-window read count in a CNA, would be a Poisson variable with both mean and variance of NDW/l. Then, the t-statistic from Equation (2) becomes

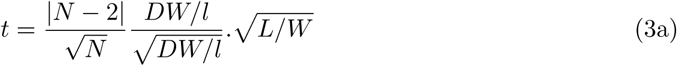

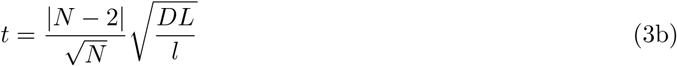

Here t is proportional to the square root of read depth D and CNA length L, and inversely proportional to the square root of the read length l.

*Remark 1*: The t statistic depends on ploidy as *|N -* 2| /N, which is a factor of 1 for N=1 or N=4, and a factor of 0.578 for N=3.

*Remark 2*: DL/l acts as a compound variable, where a doubling of D has the same effect as a doubling of L, or a halving of l. In real studies, shorter reads (smaller l) help power by increasing read counts per window, but this comes with the cost of reduced mapping accuracy, especially when l is below 50-100 nucleotides. We currently do not incorporate this reduced mappability with smaller l.

*Remark 3*: The t-statistic is not influenced by the choice of window size W in the Poisson sampling scenario. While a large W increases the mean read count in proportion to W, it increases the standard deviation by 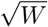 and reduces sample size by 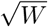, and these opposing effects cancel out each other in the expression of t. However, the *power* of the test is still indirectly dependent on W through the parameter of degrees of freedom. Only when n >> 5, i.e. with CNVs containing a large number of windows, this dependence of power on the degrees of freedom becomes negligible.

In **Figure 1**, we present an example with N=3, l=100, W=1000, and *α*=0.05, and show power as a function of CNA length L and read depth D under a Poisson model. For a CNA with L=10kb, read depth of 0.1 shows very low power; while D=1 brings much higher power. With today’s sequencing technology, 1X coverage of a mammalian genome costs $30-40. In comparison, a 3-million SNP genotyping platform will measure ∼10 SNP loci in this CNA, but at a higher cost.

**Figure 1:**
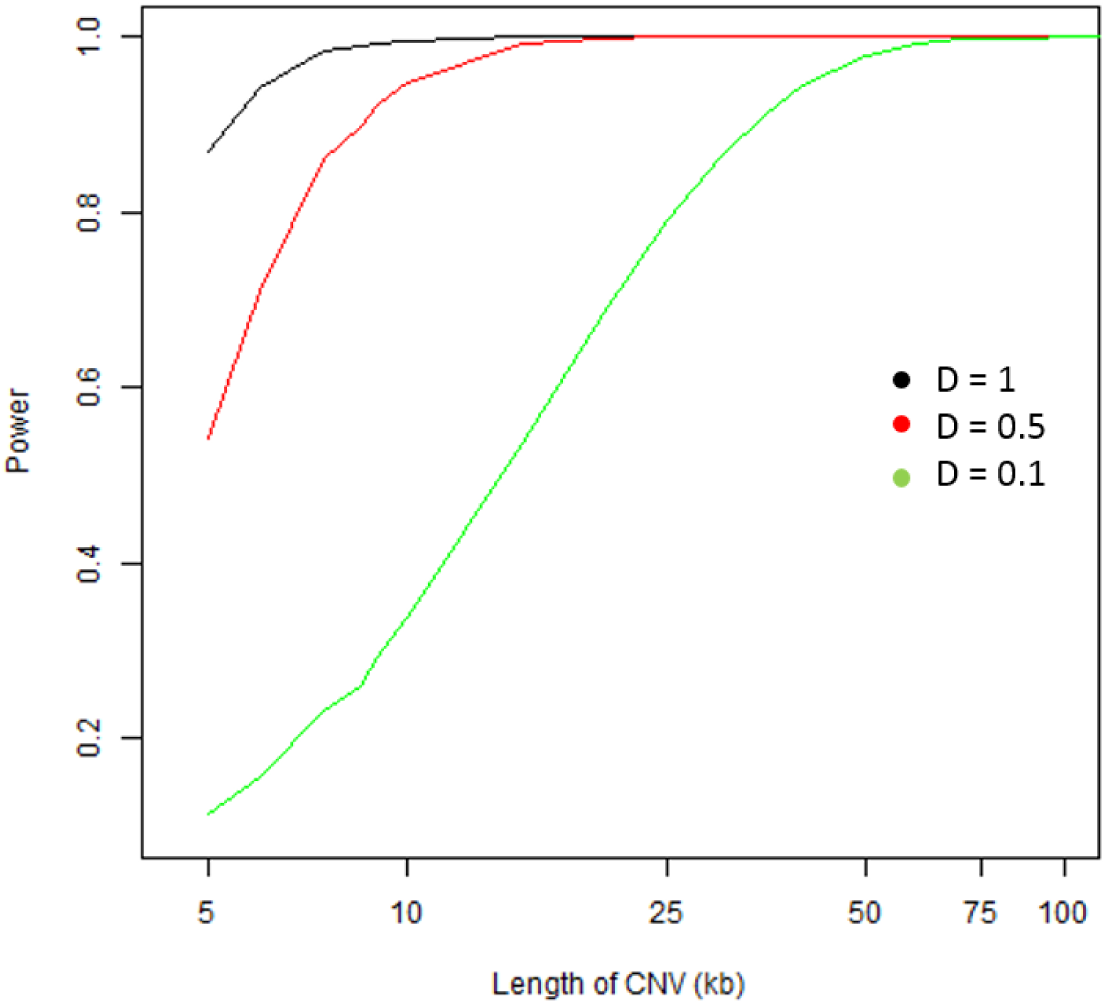
Power calculations using equation (3): Dependence of power on length of the CNV (x axis) and read depth (colors), keeping constant l=100, W=1000 and N=3.

### 4.2 Inflation of variance (Negative Binomial sampling)

Sequencing-based counting of DNA fragments along the genome may seem to mimic a Poisson sampling process; but empirical data have shown that read depth distribution exhibit an overdispersion of variance, and this is sometimes modeled as sampling from a negative binomial distribution (Anders, S and Huber, W, 2010). For example, Wang et al. (2008) showed that the variance of read depth in a whole-genome sequencing dataset was twice the average read depth. We thus consider the case where variance is inflated in the range of 1-2.

In practice, the degree of over-dispersion depends on the sequencing approach, and ultimately results from sequence-dependent molecular reactions in the library preparation process. Even with PCR free protocols some over-dispersion will remain. Further, exome-sequencing or panel-sequencing will lead to a greater dispersion than whole-genome sequencing due to the additional steps in target enrichment.

We generalize the previous section by introducing an inflation factor, *θ*, in the variance of read depth.

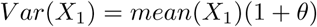

where the mean of *X*_1_ in a region of ploidy N is NWD/l as before, and *θ* is the over-dispersion parameter estimated from the sequencing data.

The term (1+*θ*) resembles the expression of variance in a negative binomial distribution, where *θ* is expressed as a variable *μϕ*, which depends on the mean *μ*. In most of what follows we examine the impact of *θ* as an independent inflation factor. However, if the negative binomial distribution is the operating model, *θ* depends on *μ*, or NDW/l. We’re in the process of making our calculator capable of modeling *θ*, modeling *ϕ*, where *θ* depends on NDW/l.*ϕ*.

The expression in Equation (3b) now has an additional term from the over-dispersion:

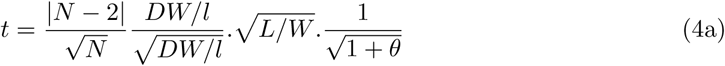

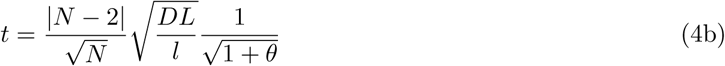

*Remark 1*: The dependence of the t statistic on D, L, l is still in the form of DL/l, as a three-way compound variable. And N’s impact is 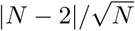, the same as before.

*Remark 2*: The inflation factor *θ* acts as a multiplier of 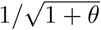. which is 0.82 for 1+*θ* of 1.5, and 0.71 for 1+*θ* of 1 (a two-fold inflation).

*Remark 3*: A change in 1+*θ* from 1 to 2 by two-fold is equivalent to a 2-fold change in DL/l. Thus (1+*θ*) act with similar impact as D, L and the inverse of l.

Keeping the same parameters as in Figure 1, with N=3, l=100, W=1000 and *α*=0.05, we estimated the effect of *θ* on power as it varies with D and L (**Figure 2**). As predicted, a 2-fold decrease of read depth from 1 to 0.5 is equivalent to a two-fold decrease of 1+*θ* (the black dashed line is identical to the red solid line).

**Figure 2:**
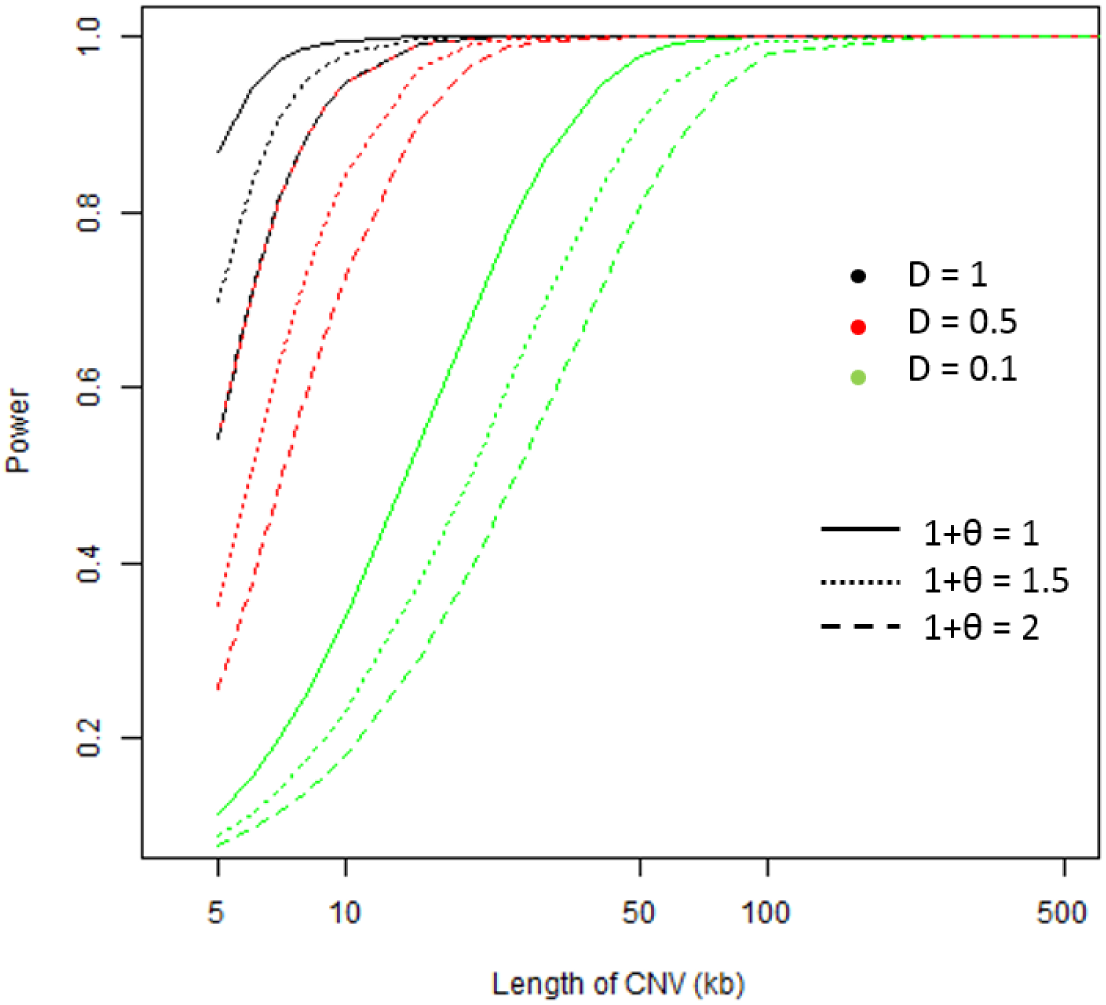
Power calculations using using equation (4): Dependence of power on length of the CNV (x axis), read depth (colors) and *θ* (line types), keeping constant l=100, W=1000 and N=3, as in Figure 1.

## 5 Two-sample mixing (F < 1)

In cases of two-sample mixing, we denote the fraction of the CNA-bearing population as F. The DNA dosage (“ploidy”) in the CNA region is a weighted average: NF+2(1-F). Thus, the per-window read count in the CNA is a random variable with a mean of (F(NDW/l) + (1-F)(2DW/l)).

From Equation 2, the t statistic is:

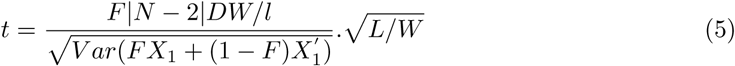

where *X*_1_ is the read counts derived from the N-ploid population at fraction F and a mean of NDW/l, and 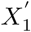 is the read count from the diploid population at fraction (1-F) and a mean of 2DW/l.

When the CNA-bearing fraction is very low, as in the case of low tumor cell fraction in circulating blood, F ≪ 1, and by omitting the *FX*_1_ term in the denominator we have

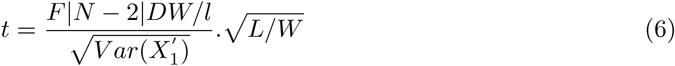

### 5.1 No variance inflation (Poisson sampling)

Under the assumptions of Poisson sampling, variance equals the mean and the denominator in Equation (5) becomes:

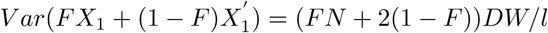

Thus,

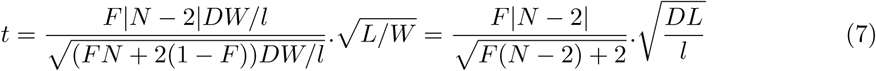

*Remark*: Here F affects both the numerator and denominator. The effect size in the numerator scales linearly with F; whereas the standard deviation in the denominator is the square root of F(N-2)+2. The relationship among the other parameters stays the same as in Equation (3b).

**Figure 3** extends the earlier example where N=3, l=100, W=1000 and *α*=0.05 by showing the reduction of power when F=0.5 and 0.1. The effect of a 2-fold change in F is far greater than a two-fold change of D, and this is because the effect of F is nearly linear with F, yet the effect of D, L, or 1/l is linear with their square root value. In other words, a two-fold reduction of purity requires nearly 4-fold larger sequencing depth or longer length of CNA to reach equivalent power.

**Figure 3:**
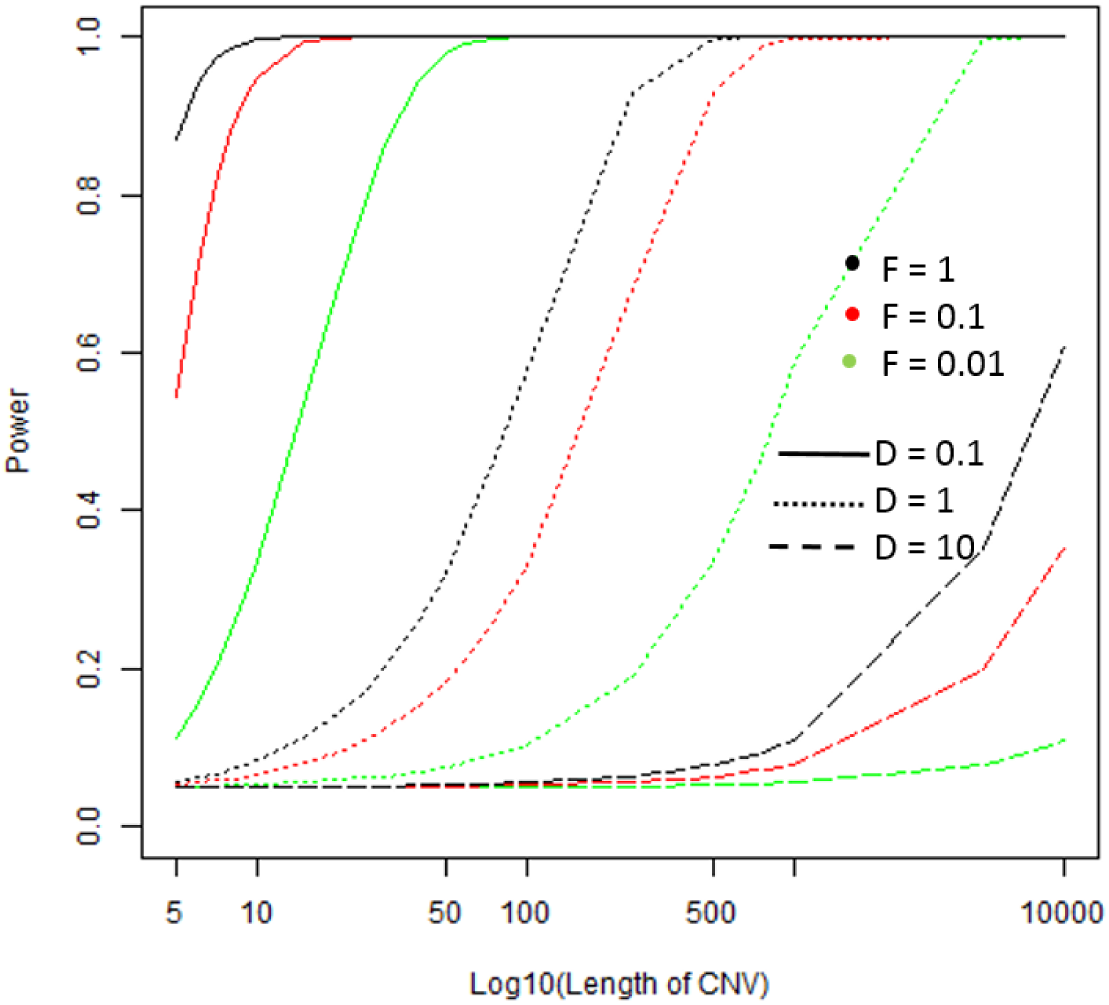
Power calculations using equation (7): Dependence of power on length of the CNV (x axis), read depth (colors), and tumor fraction (line type), keeping constant l=100, W=1000 and N=3, as in Figure 1.

If we assume low sample purity (F ≪ 1), the denominator in Equation (6) becomes: 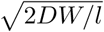, And Equation (7) simplifies to:

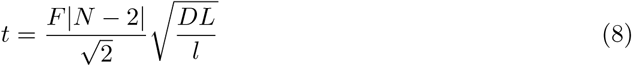

Here, F only acts on the numerator, and its effect is equivalent to the effect of the square root of the composite variable (DL/l).

### 5.2 Inflation of variance (Negative Binomial sampling)

As in 4.2 we add an inflation term (1+*θ*) to the denominator in equation (5). Here *θ* can be considered either as an independent variable or as *μϕ*, where *μ* is the mean and *ϕ* is an independent variable. When *θ* takes on the latter form, *μϕ*, the variance is from a negative binomial distribution, where the inflation factor is a function of NDW/l. In our calculator we allow both the *θ* and the *μϕ* formulations.

When Eq.5 and 7 are extended by adding (1+*θ*), it becomes

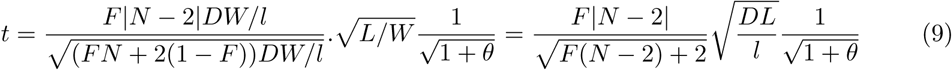

**Figure 4** shows the dependence of power on read depth D, length of the CNV L, and over-dispersion *θ* using this model, where we fix N=3, l=100, W=1000 and *α*=0.05.

**Figure 4:**
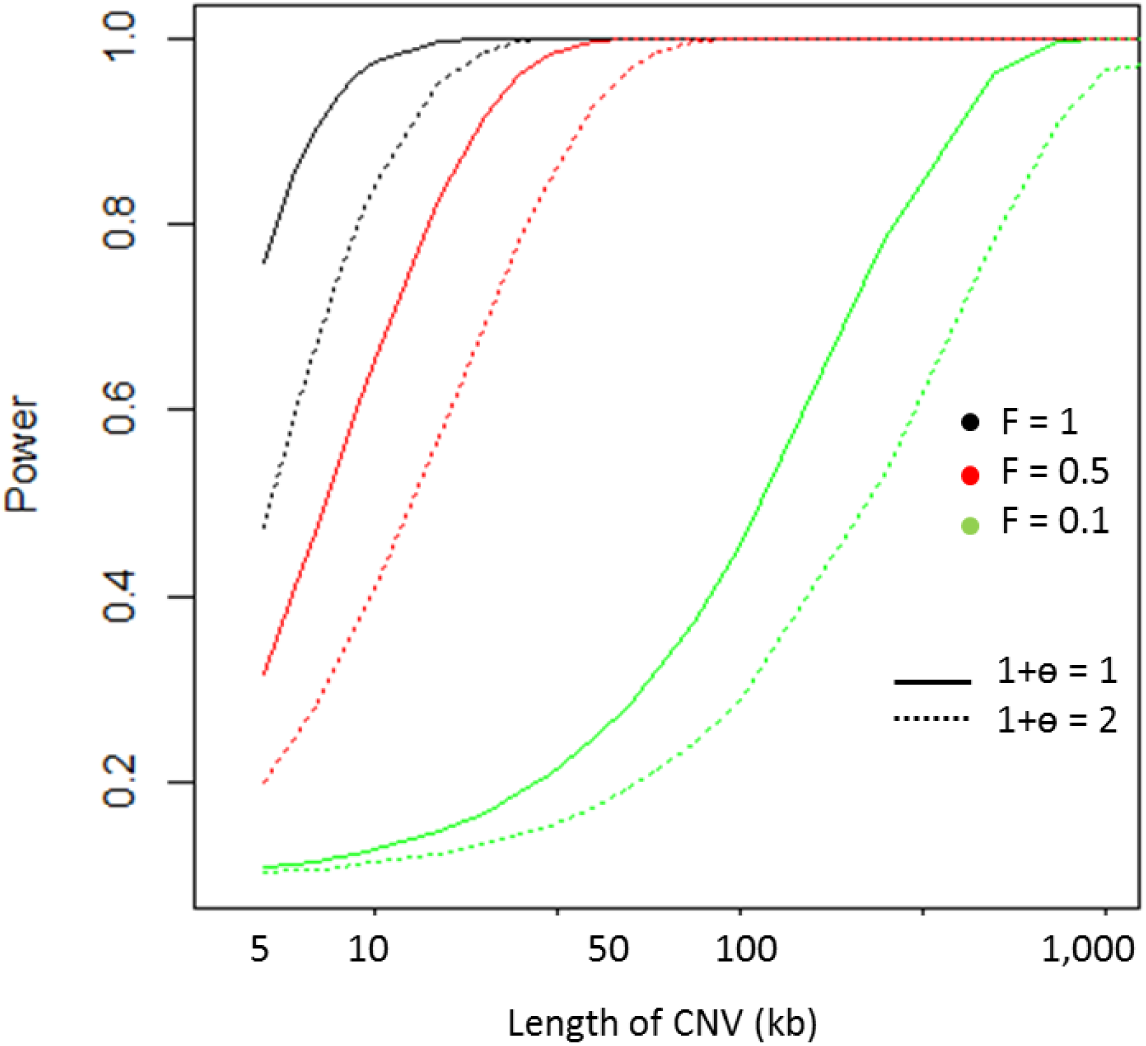
Power calculations using equation (9): Dependence of power on length of the CNV (x axis), tumor fraction (colors), and (1 + *θ*)(line type), keeping constant D=1, l=100, W=1000 and N=3.

Under assumptions of very low sample purity (F ≪ 1), equation (9) becomes:

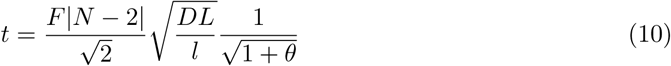

## 6 Online Calculator

We present an online calculator at https://shiny.med.umich.edu/apps/hanyou/CNV_Detection_Power_Calculator/ implementing these formulae, in which users can design experiments optimized to their experimental parameters.

## 7 Use Cases

### 7.1 Large CNV, ultra-shallow sequencing

When sequencing tumor-derived DNA, sometimes one faces the situation of low budget or low amounts of tissue material, or one wishes to use shallow sequencing to gain a preliminary glimpse of the tumor-normal mixing ratio. These situations share the common constraint of small D; and the power analysis aims to answer questions like these:

Q1. If coverage D is in the range of 0.1-1, and purity F is in the range of 0.01-1, how large L needs to be to have adequate power to be found?

Q2. How much should one worry about over-dispersion (1+*θ*)?

While the situation of F=1 has been shown in Figure 1, we used our calculator to demonstrate (**Figure 5**) how power decreases with F in (0.5, 0.1, 0.01) and with (1+*θ*) increases from 1 to 2. As expected, (1+*θ*) has a relatively small impact, whereas F has a much larger impact than D. At D=1 and F=1, a 5 kb CNA has 80% power to be detected (**Figure 1**). However, when F drops to 0.5, 0.1, and 0.01, 80% power is only achieved for much longer CNA, at L of ∼10 Kb, >100 Kb, and >10 Mb, respectively.

**Figure 5:**
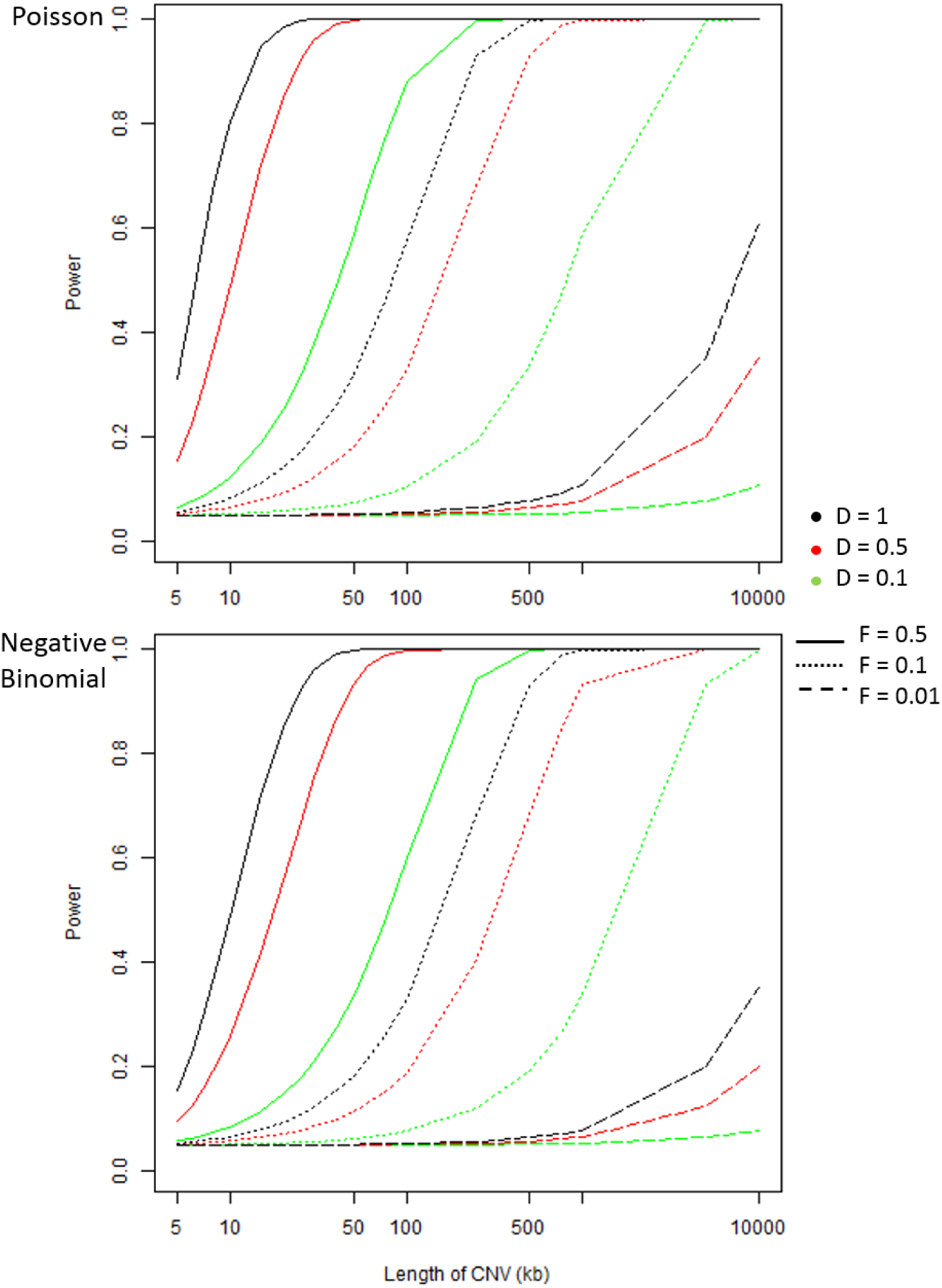
Use Case 1: Dependence of power on length of the CNV (x axis), read depth D (colors) and tumor fraction F (line types) keeping constant l=100, W=1000 and N=3. Poisson model is shown in the top panel and the negative binomial model (with *θ*=1) on the bottom.

### 7.2 Ultra-deep sequencing, low purity

Sometimes the researcher already knows the regions of the genome to focus on, and is interested in detecting very rare cell populations with copy number aberrations in these regions, even if it requires ultra-deep targeted sequencing. Such needs arise in (1) monitoring of potential recurrence of a particular cancer clone during remission and hoping to achieve high sensitivity with a large D; (2) monitoring gene-specific aberrations in circulating tumor cells or circulating tumor DNA when a predefined list of oncogenes are particularly relevant for a particular cancer type (such as BRAF in melanoma). These practical demands share the common feature of small F and large D, with questions such as these:

Q1. If the known CNA covers n1 exomes, where n1 is the sample size, akin to a fixed L/W, how deep (D) do I need to sequence to have high power to detect the remission when this clone rises to F=1%?

Q2. If I can design a new panel of n1 capture probes (or amplicons) to cover the known CNA, how wide (n1) and how deep (D) do I need to sequence to achieve optimal sensitivity given a fixed cost, which contains both probe design cost and sequencing cost?

Q3. If I have a platform with known n1, but a fixed annual budget, should I sequence the time series samples shallowly (smaller D) but more frequently, or deeply but less frequently? How can I translate this question into the relative clinical value of detecting a clone at F1 through weekly surveillance, vis-a-vis detecting this clone at an even lower frequency F2<F1 but through monthly surveillance, hence a possibly delayed detection?

Our calculator allows the user to explore the joint effects of all the relevant parameters. As an illustrative example, in **Figure 6** we plot the power in the range of low purity: F in the range of 0 −0.1, and ultra-deep sequencing: D in the range of 1 - 1000. A 5-10 Kb CNA can be detected at F of 2% if D is as high as 1000×. But the power drops to 10-30% when D is only 100×.

**Figure 6:**
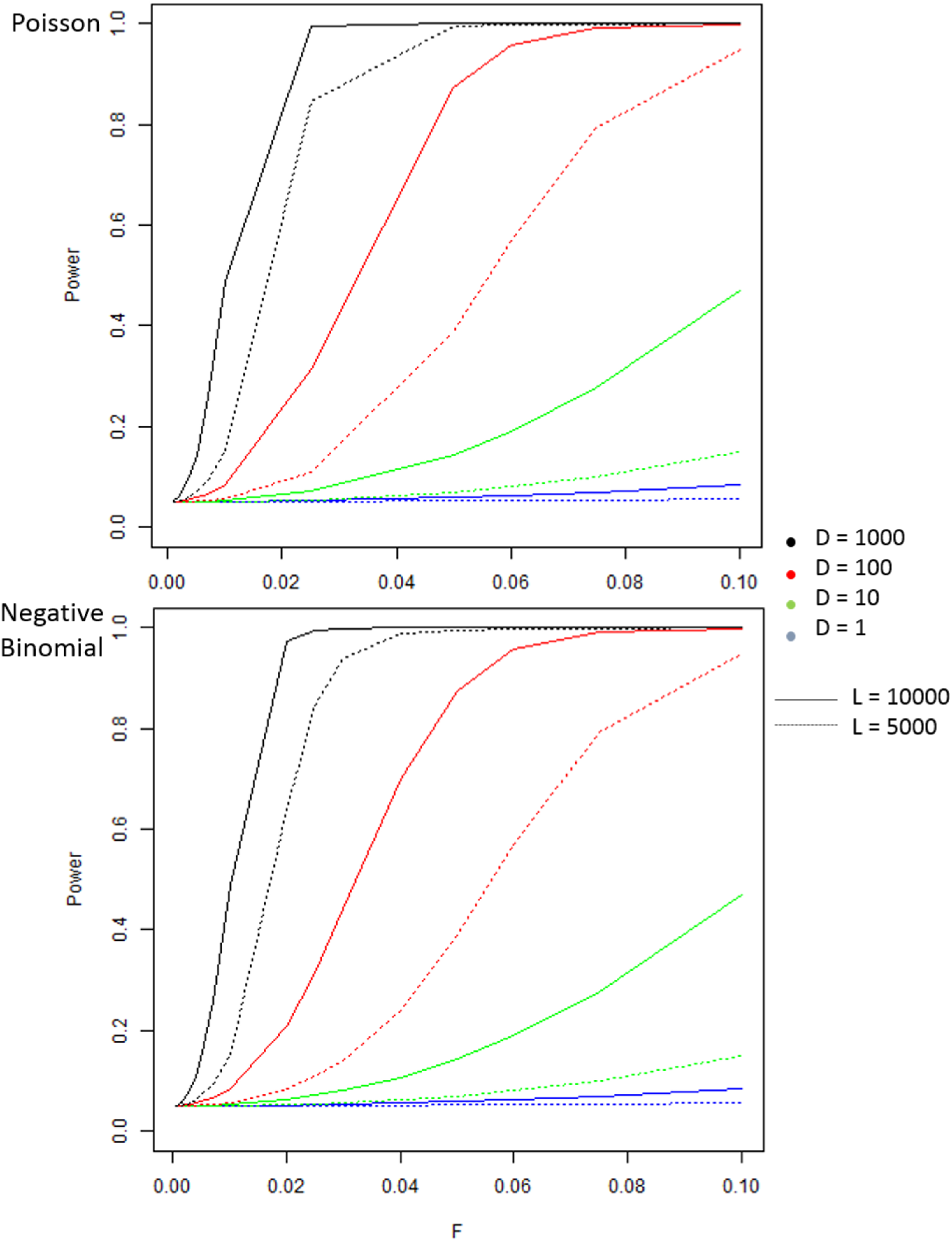
Use Case 2: Dependence of power on tumor fraction (x axis) and read depth D (different colors) keeping constant l=100, W=1000, N=3. Poisson model is shown in the top panel and the negative binomial model on the bottom (with *θ* of 1)

### 7.3 Parameter estimation: sample purity F and its confidence interval

In some situations, the researcher may wish to report the mixing ratio F for a given CNV, along with an estimated confidence interval. Our framework can be easily turned from hypothesis testing to effect estimation. Testing *H*_*A*_: *t* ≠ 0 is equivalent to testing the compound hypothesis *F* ≠0 & *N ≠*2. Here we focus on estimating F.

The per-window read count in the CNA is a random variable with an estimated value of:

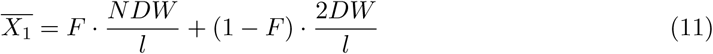

From this expression, we can derive F as:

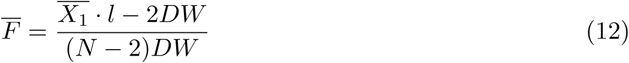

The standard deviation of F can then be expressed as:

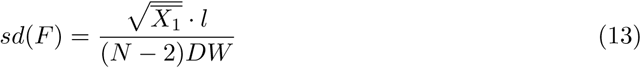

Hence the confidence interval of F at significance level *α* is expressed as:

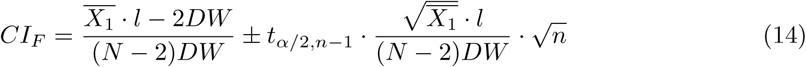

where n=L/W is the number of windows in the CNA. When n is large, *t*_*α/*2, *n* − 1_ can be approximated by the standard normal variable. For *α* = 0.05, *Z*_*α/*2_ is 1.96.

## 8 Discussion

In this study we systematically evaluated the factors affecting statistical power to detect CNAs from sequencing read depth data. We show that in a basic Poisson-sampling model the overall signal for a stepwise change of DNA ploidy can be expressed as a t score, and it depends on ploidy N by 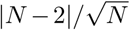, and is affected by CNA length L, read length l and sequencing depth D by *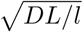*. The width of the genomic window defined by the analyst to bin the sequence reads, W, has opposing effects on three components of the t score and they cancel out, thus the choice of W has no impact on power, so long as the CNA contains more than a few windows to allow a high degrees of freedom for the t score.

We then explored two complications, summarized in **Table 1**. First, if the variance of read counts along the genome is more than the mean, we introduce an inflation factor of (1 + *θ*) to the variance to reflect its over-dispersion beyond the Poisson sampling. Our calculator either models *θ* as an independent parameter, or as *μϕ*, which represents the negative binomial model. In the former case, t is modified by a multiplying term 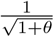. And in the latter case, *θ* is expanded as the product of the independent parameter *ϕ* and the mean read count in the N-ploid region *μ*=NDW/l.

**Table 1.**
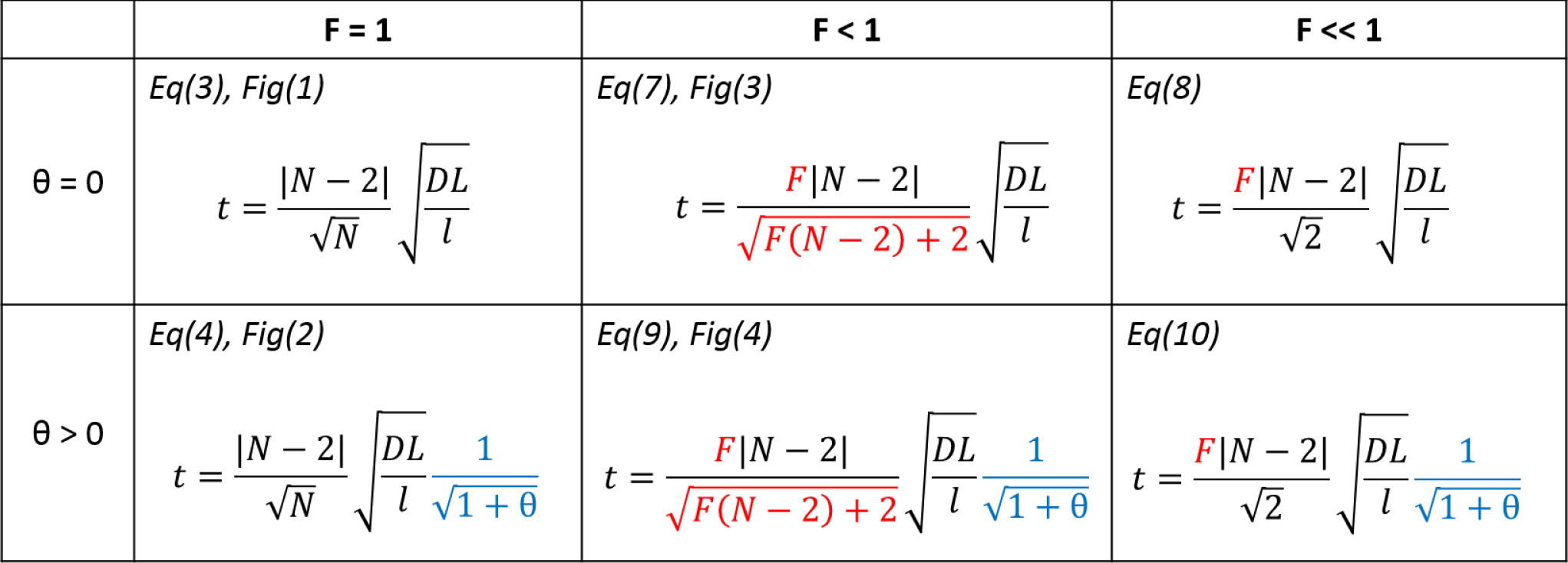
The t statistic for each model we estimate. The complexity of the model increases from left to right (due to tumor purity) and from top to bottom (due to addition of the variance inflation factor.)

In the second complication, we consider two-sample mixing where the CNA-bearing fraction is F, also known as “purity” in cases of tumor-normal mixing. F’s impact on the t score is entwined with N, as in 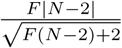. When F=1, this term becomes 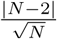, as in the simplest case. When F ≪ 1, this term becomes 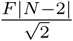, where the t score is a linear function of F and N. Importantly, in most cases t depends on F but only the square root of D, L, l, and 1 + *θ*. Thus tumor purity has a largest impact: a two-fold change in F can only be compensated by a four-fold change to the opposite direction by D, L, 1/l, or 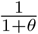.

This analytical framework relies on several simplifying assumptions. While accounting for an inflation of the variance of read counts, we modeled this inflation either as (1 + *θ*) or as (1 + *μϕ*). However real-life data may be more complex, and certainly platform-dependent. New sequencing methods become available constantly, and targeted-sequencing approaches such as exome sequencing or targeted gene-panel sequencing brings uneven distribution of the sampled regions where sample size is no longer simply L/W, rather it is determined by the empirical distribution of the targeted regions. In such cases, our model can be generalized by using n1 (the number of regions) rather than L/W as the parameter for sample size, allowing the user to calculate power with n1 as the known number of observations in the CNA. Whole-exome sequencing is also subject to variation due to capture efficiency of individual probes; and this may vary from one experiment to the next. Users of our calculator need to exercise caution when dealing with capture rate heterogeneity or otherwise biased sampling processes.

While we illustrated the influence of 1 + *θ* in the range of 1-2, in practice this factor can be bigger if regional biases have not been adequately adjusted; but conversely, it can be significantly smaller if we increase window size W. That is, if the variance of read depth is over-dispersed above the mean by 1 + *θ*, after binning reads and tallying the per-window read counts, the variance of the aggregated counts is inflated by a factor far smaller than 1 + *θ* (simulation results, not shown). Thus in practice, if the CNA is sufficiently large and one can choose a large W, the inflation factor 1 + *θ* may be very close to 1 and a Poisson model may be adequate.

A major limitation of our analysis is that it only utilized the information from read depth patterns, while there have been other approaches that also take into account split reads (Jiang et al., 2012), paired-end information (Korbel et al., 2009; Gillet-Markowska et al., 2015), single nucleotide polymorphisms, RNA-Seq data, or a combination of these signal sources (Hormozdiari et al., 2009; Handsaker et al., 2015; Layer et al., 2014). Incorporating these complementary types of information would increase the power of CNV detection.

Another major limitation is that read depth data depend on the quality of the alignment, which can be compromised for a species without a high-quality reference genome, or for repetitive regions in general. Sometimes corrections can be made if systematic bias is learned from sequencing data from many samples, and the adoption of long-read sequencing or linked-read assembly is expected to resolve the most difficult repetitive regions.

